# Cytokines in the Urine of AKI patients regulate TP53 and SIRT1 and can be used as biomarkers for the early detection of AKI

**DOI:** 10.1101/2023.01.20.524775

**Authors:** Lars Erichsen, Chantelle Thimm, Wasco Wruck, Daniela Kaierle, Manon Schless, Laura Huthmann, Thomas Dimski, Detlef Kindgen-Milles, Timo Brandenburger, James Adjaye

## Abstract

Acute kidney injury (AKI) is a major kidney disease with a poor clinical outcome. It is a common complication with an incidence of 10-15% of patients admitted to hospital. This rate even increases for patients who are admitted to the intensive care unit with an incidence of >50%. AKI is characterized by a rapid increase in serum creatinine, decrease in urine output, or both. Associated Symptoms include feeling sick or being sick, diarrhoea, dehydration, decreased urine output-although occasionally urine output remains normal, fluid retention-causing swelling in the legs or ankles, shortness of breath, fatigue and nausea. However, sometimes acute kidney injury causes no signs or symptoms and is detected by lab tests. Therefore, an urgent demand for non-invasive biomarkers for early detection of AKI are highly desirable. This might enable the prevention of the progression from AKI to CKD.

In this study, we analysed the secretome of urine of an AKI patient cohort employing a kidney-biomarker cytokine assay. Based on these results we suggest, ADIPOQ, EGF and SERPIN3A as potential biomarkers, which might be able to detect AKI as soon as 24 h post-surgery. For the later stages, common biomarkers for the detection of AKI in both male and female patients we suggest, VEGF, SERPIN3A, TNFSF12, ANPEP, CXCL1, REN, CLU and PLAU. These markers in combination might present a robust strategy to identify the development of AKI as early as 24h or 72h post-surgery. Furthermore, we evaluated the effect of patient and healthy urine on human podocyte cells. We conclude that cytokines in the urine of AKI patients trigger processes which are needed to repair the damaged nephron and activate TP53 and SIRT1 to maintain the balance between proliferation, angiogenesis, and cell cycle arrest. In conclusion, the Renin-Angiotensin pathway seems to have major implications.

## Introduction

The understanding of the ethology underlying AKI has fundamentally changed in recent years^1^. In the past, research was focused on the most severe impairment of the kidney. More recent studies have provided evidence that even minor injuries or impairment of kidney function can lead to severe consequences such as changes in serum creatinine (sCr) and/or urine secretion (UO)^2^. Today the term acute kidney injury (AKI) is used for a major kidney disease with a poor clinical outcome. It is characterised as an abrupt (within hours or days) decline in renal function, which includes structural damage to the nephron as well as loss of functionality. It is a common complication for patients admitted to the hospital with an incidence of 10-15% ^3^. This rate even increases for patients who are administered to the intensive care unit with an incidence rising to >50% ^4^. The classification of AKI includes pre-renal AKI, acute post-renal obstructive nephropathy, and intrinsic acute kidney disease. Of these, only the latter represents a true renal disease, whereas pre- and post-renal AKI are the result of extra-renal disease leading to decreased glomerular filtration rate (GFR). If these pre- and/or post-renal conditions persist, they eventually develop into cellular damage to the kidney and thus intrinsic renal disease^2^. The current diagnostic approach for AKI is based on an acute decrease in GFR reflected by an acute increase in sCr level and/or decrease in UO value over a given time interval^2^. According to the current kidney disease improving global outcomes (KDIGO), the definition AKI is classified into three distinct stages: Stage 1: Creatinine > 1.5 times baseline or increase of ≥0.3 mg/dL within any 48 h period, or urine volume <0.5ml/kg for 6 – 12 h. Stage 2: Creatinine ≥2.0 times baseline or urine volume <0.5 ml/kg for ≥12 h and Stage 3: Creatinine >3.0 times baseline or increase to ≥4.0mg/dL or acute dialysis, or urine volume <0.3 ml/kg for ≥24 h^5^. If these conditions persist for up to seven days, the pathological conditions are referred to as AKI. Between 7 and 90 days, the conditions are referred to as acute kidney disease (AKD) and if they exceed 90 days chronic kidney disease (CKD) has manifested^6^. In their clinical presentation, kidney disease are very difficult to identify, because except for urinary tract obstructions, they do not cause any specific signs or symptoms^6^. Furthermore, diagnosis of AKI is hindered by the fact that specific syndromes often co-exist (as reviewed in ^6^) or AKI arises as part of other syndromes such as heart or liver failure or sepsis. In addition 40% of all AKI cases in the hospital are related to surgical procedures^7^ with AKI being associated with cardiac surgery with an incidence ranging from 7-40%^8–11^.

Therefore, an urgent demand for non-invasive biomarkers for early detection of AKI are highly desirable. This might improve patient outcome in the intensive care unit and reduce the risk of progression from AKI to CKD and ultimately End Stage Kidney Disease-ESKD

Under healthy physiological conditions kidney cells divide at very low rates^12,13^. In contrast, several studies have reported a dramatic increase in proliferating cells after AKI, which can also be accompanied by transient dedifferentiation^13,14^ to repair the injured nephron. This repair process has been identified to be possible maladaptive depending on the severity of the injury, involving increased DNA damage and cellular senescence^15^. Therefore, it is not surprising that TP53 and its downstream target TP21 were found to be upregulated in the kidney after AKI and inhibition or gene deletion reduces kidney lesions^16–20^. Interestingly, using P53 inhibitors, after unilateral ischemia reperfusion injury resulted in reduced fibrosis^19^. Despite this progress in understanding the role of TP53 in the pathology of kidney injury, the underlying mechanism in the progress remains largely unknown. The NAD+-dependent class III histone deacetylase SIRT1 is involved in many biological processes including DNA damage repair and TP53 activation^21,22^. Furthermore, SIRT1 has been reported to play a protective role in many diseases including AKI^23–25^.

In the present study, we screened urine from intensive care patients who developed AKI 24 h and 72 h post-surgery for known kidney biomarkers employing a kidney-injury cytokine assay. We identified ADIPOQ, EGF and SERPIN3A as potential biomarkers capable of detecting AKI as early as 24h post-surgery. At 72 hrs post-surgery, common biomarkers for the detection of AKI in both male female patients include, VEGF, SERPIN3A, TNFSF12, ANPEP, CXCL1, REN, CLU and PLAU. Furthermore, we applied the urine to our recently published hTERT immortalized podocyte cell line^26^ and evaluated the effects of the urine on TP53 and SIRT1 expression.

## Results

### Identification of AKI biomarkers in the Urine of patients 24h post operation

Urine samples from six stage 2/3 AKI patients and six patients without AKI were collected 24 h post-surgery at the clinic for anesthesiology at Heinrich Heine University-Düsseldorf. The patients without AKI were aged between 55 and 75 years-three males and three females. The AKI patients were aged between 34 and 84 years with the same sex distribution (see table 1-Material and Method section). The urine samples were incubated individually on the Human kidney biomarker Array, scanned images are shown in supplementary figure 1. The results are shown in figure 1. As shown in the dendrogram (fig. 1a) the samples segregated into two distinct clusters of AKI (red) and healthy controls (blue). The clustering of the AKI group could even further be sub-divided into male and female. Interestingly, cytokines in the healthy urine seem to be more equally distributed between the genders according to the dendrogram. The heatmap (fig. 1b) and the barplots (fig. 1c and 1d) show cytokines found to be significantly regulated as detected by the array.

**Table 1:** Patient cohort.

| Parameter | Total (n=12) | No AKI (n=6) | AKI 2 and 3 (n=6) |
| --- | --- | --- | --- |
| **Age [years]** | 68 ± 13 | 68 ± 8 | 69 ± 18 |
| **Gender** |  |  |  |
| Female [%] | 6 (50) | 3 (50) | 3 (50) |
| Male [%] | 6 (50) | 3 (50) | 3 (50) |
| **Comorbidities** |  |  |  |
| Obesity [%] | 5 (42) | 2 (33) | 3 (50) |
| Hypertension [%] | 9 (75) | 5 (93) | 4 (67) |
| Diabetes Mellitus [%] | 4 (33) | 2 (33) | 2 (33) |
| CKD [%] | 2 (17) | 0 (0) | 2 (33) |
| **Preoperative Creatinine [mg/dl]** | 1,52 ± 0,76 | 1,18 ± 0,44 | 1,85 ± 0,90 |
| **Operation time [minutes]** | 284 ± 73 | 282 ± 66 | 287 ± 86 |
| **CPB time [minutes]** | 180 ± 57 | 191 ± 65 | 170 ± 51 |
| **X-clamp [minutes]** | 106 ± 42 | 109 ± 33 | 103 ± 52 |
| **Creatinine postop. [mg/dl]** | 1,33 ± 0,38 | 1,2 ± 0,34 | 1,43 ± 0,42 |
| **Creatinine on day 3 [mg/dl]** | 1,60 ± 1,08 | 1,08 ± 0,50 | 2,03 ± 1,29 |
| **Lactate postop. [mmol/l]** | 5,0 ± 3,5 | 2,5 ± 0,9 | 7,5 ± 3,2 |
| **Lactate on day 3 [mmol/l]** | 1,96 ± 1,75 | 2,2 ± 2,5 | 1,7 ± 0,5 |
| **APACHE-II postop.** | 29 ± 4 | 26 ± 2 | 32 ± 3 |
| **Ventilation [hours]** | 103 ± 192 | 20 ± 8 | 186 ± 253 |
| **LOICUS [days]** | 8 ± 10 | 2 ± 2 | 13 ± 11 |
| **LOHS [days]** | 30 ± 20 | 18 ± 12 | 42 ± 20 |
| **Last Creatinine [mg/dl]** | 1,55 ± 0,76 | 1,03 ± 0,33 | 2,07 ± 0,72 |
| **Mortality in hospital** | 1 (8) | 0 (0) | 1 (17) |

Data are expressed as mean ± SEM or number (percentage).
CPB: cardiopulmonary bypass; LOICUS= length of intensive care unit stay; LOHS = length of hospital stay

**Figure 1:**
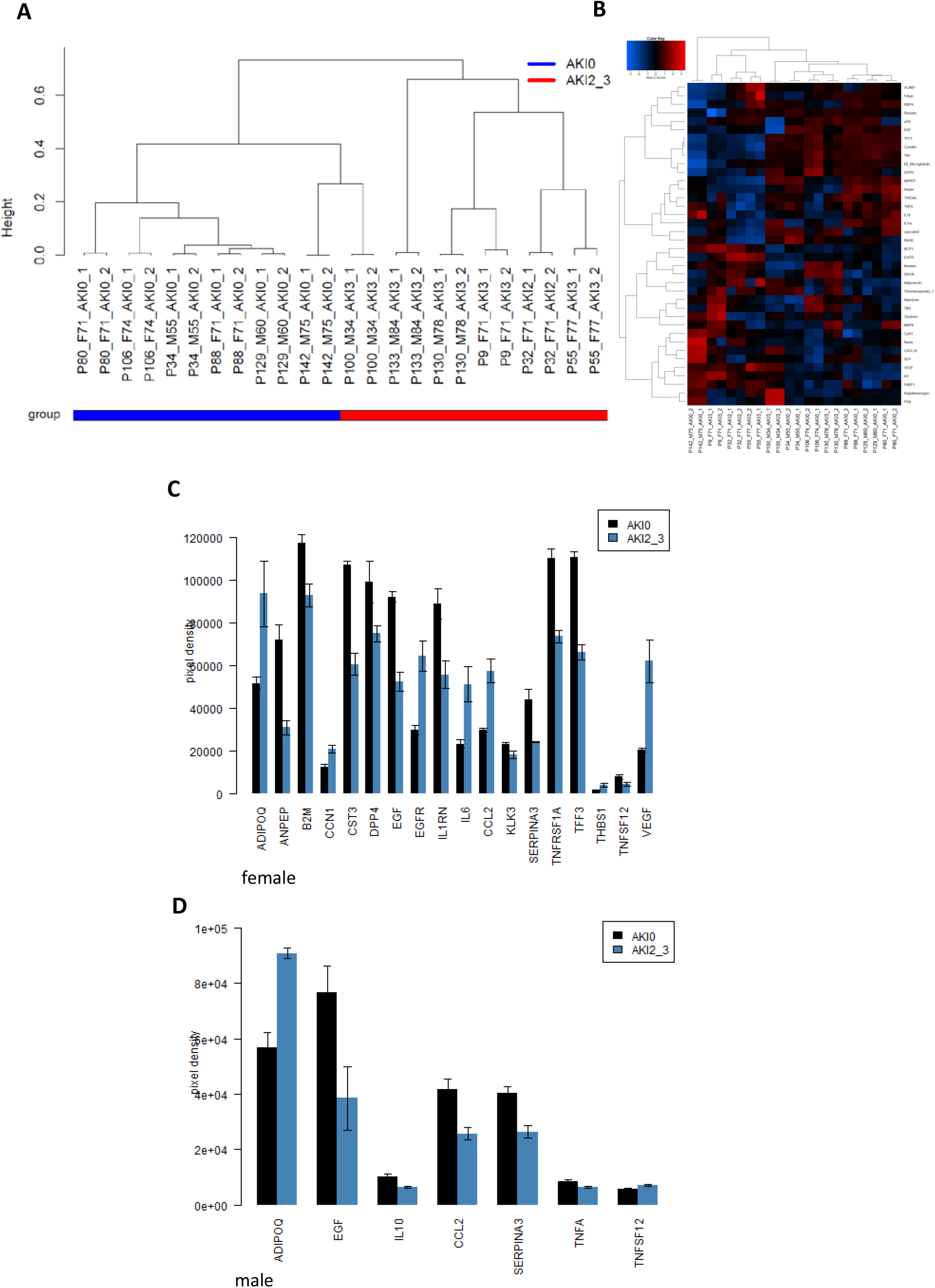
AKI biomarkers identified in patients 24 h post-surgery. (A) experiments cluster into CKD and healthy control based on the global kidney cytokine expression. (B) Heatmap and (C) barplot of markers in male (M) and barplot of markers in female (D).

In female patients, Adiponectin (ADIPOQ), Cellular Communication Network Factor 1 (CCN1), Epidermal growth factor receptor (EGFR), Interleukin 6 (IL6), CC-chemokine ligand 2 (CCL2), Thrombospondin 1 (THBS1) and Vascular Endothelial Growth Factor (VEGF) were found to be upregulated while Alanyl Aminopeptidase (ANPEP), β2-Mikroglobulin (B2M), Cystatin C (CST3), Dipeptidylpeptidase-4 (DDP4), Epidermal growth factor (EGF), Interleukin 1 Receptor Antagonist (IL1RN), Kallikrein Related Peptidase 3 (KLK3), Serpin Family A Member 3 (SERPIN3A), Tumour necrosis factor receptor superfamily member 1A (TNFRSF1A), Trefoil Factor 3 (TFF3) and Tumour necrosis factor ligand superfamily member 12 (TNFSF12) were found to be downregulated.

In male patients, Adiponectin (ADIPOQ) and Tumour necrosis factor ligand superfamily member 12 (TNFSF12) were found to be upregulated, while Epidermal growth factor (EGF), Interleukin 10 (IL10), CC-chemokine ligand 2 (CCL2), Serpin Family A Member 3 (SERPIN3A) and Tumour necrosis factor alpha (TNFA) were found to be downregulated.

### Identification of AKI biomarkers in the Urine of patients 72h post operation

Urine samples from the same six AKI and non-AKI patients were collected 72h post-surgery. Patient characteristics are similar between AKI and non-AKI patients (e.g., age, gender, comorbidities, table. 1). Intra-operative data (operation time, cross clamp time and time on cardiopulmonary bypass) are also comparable. Creatinine levels as expected are higher in the AKI group, and so are the lengths of intensive care and hospital stay. The urine samples were pooled (based on gender and disease state) and incubated on the Human kidney biomarker Array-results are shown in figure 2, -scanned images are shown in supplementary figure 2. As shown in the dendrogram (fig. 2a) the samples segregated into two distinct clusters with the male AKI samples (black) being closer but distinct from the healthy controls (red), compared to the female AKI samples (blue). The heatmap (fig. 2b) and the barplots (fig. 2c and 2d) show cytokines significantly regulated (p < 0.05) as detected by the array. In female patients, Alanyl Aminopeptidase (ANPEP), Chemokine (C-X-C Motif) Ligand 1 (CXCL1), Interleukin 1 Receptor Antagonist (IL1RN), Renin (REN) and Vascular Endothelial Growth Factor (VEGF) were found to be upregulated, while β2-Mikroglobulin (B2M), Clusterin (CLU), Epidermal growth factor (EGF), CC-chemokine ligand 2 (CCL2), Matrix metallopeptidase 9 (MMP-9), Membrane Metalloendopeptidase (MME), Kallikrein Related Peptidase 3 (KLK3), Advanced Glycosylation End-Product Specific Receptor (AGER), Serpin Family A Member 3 (SERPIN3A), Tumour necrosis factor ligand superfamily member 12 (TNFSF12) and Plasminogen Activator, Urokinase (PLAU) were found to be downregulated. In male patients, Alanyl Aminopeptidase (ANPEP), Epidermal growth factor receptor (EGFR), Alpha 2-HS Glycoprotein (AHSG), Chemokine (C-X-C Motif) Ligand 1 (CXCL1), CC-chemokine ligand 2 (CCL2), Renin (REN) and Vascular Endothelial Growth Factor (VEGF) were found to be upregulated, while Annexin A5 (ANXA5), Clusterin (CLU), Cellular Communication Network Factor 1 (CCN1), Interleukin 1 Receptor Antagonist (IL1RN), Interleukin 6 (IL6), Interleukin 10 (IL10), SKP1-CUL1-F-box protein (SKP1), Serpin Family A Member 3 (SERPIN3A), Tumour necrosis factor ligand superfamily member 12 (TNFSF12) and Plasminogen Activator Urokinase (PLAU) were found to be downregulated.

**Figure 2:**
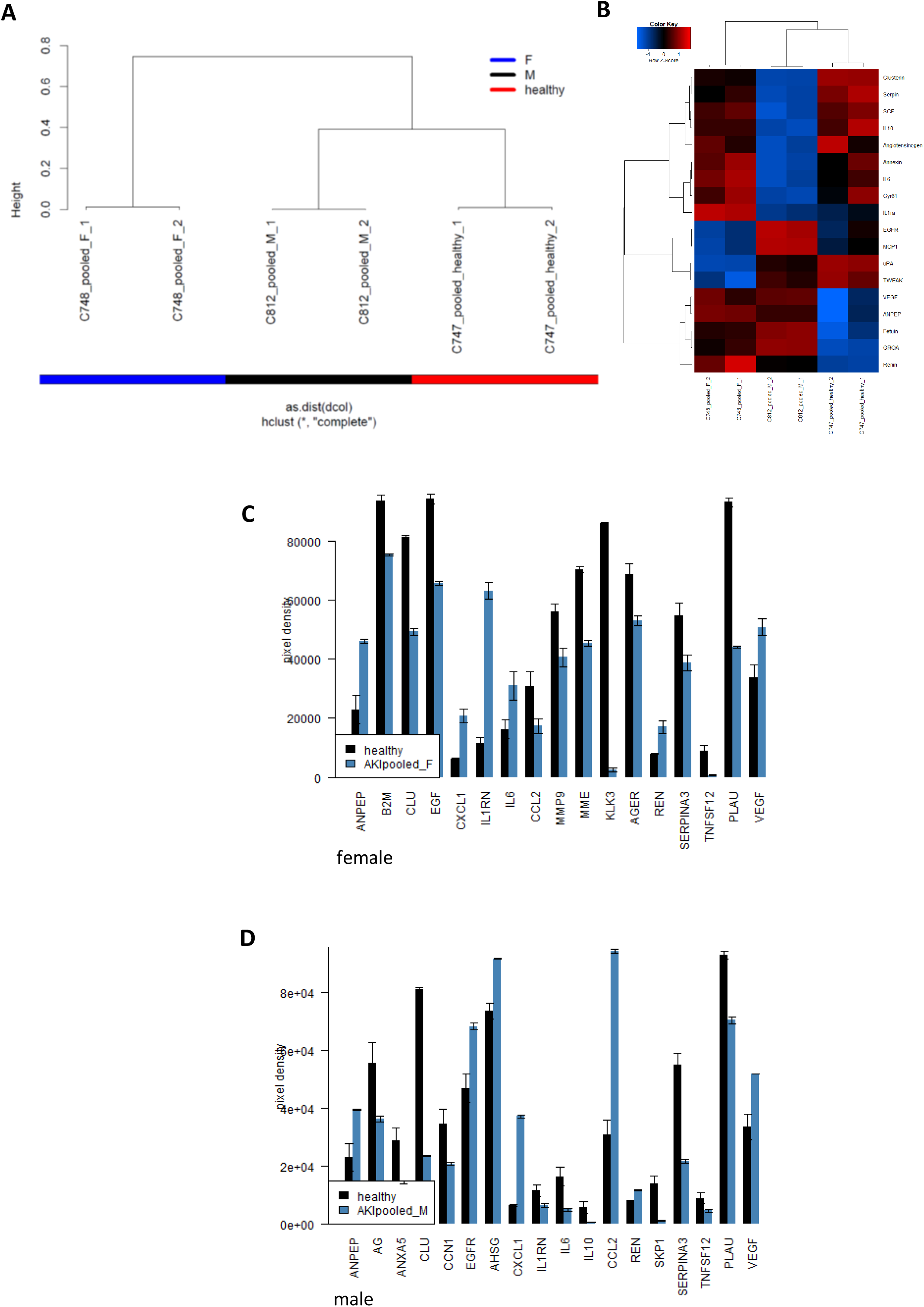
AKI biomarkers identified in patients 72 h post-surgery. (A) experiments cluster into CKD and healthy control based on the global kidney cytokine expression. (B)Heatmap and (C) barplot of markers in male (M) and barplot of markers in female (D).

### Effect of AKI stage 2/3 urine on podocytes

A number of the chemokines, such as VEGF or EGFR, that were found to be upregulated are capable of either activating or being activated by either TP53^27–30^ or SIRT1^31,32^. To test the hypothesis that the chemokines found to be upregulated in the AKI stage2/3 urine have a direct influence on the expression levels of TP53 and SIRT1, we incubated our recently published immortalized podocyte cell line UM51 hTERT^26^ with medium composed of 10% of the patient urine 72hrs post-surgery for 120 h. Relative protein expression normalized to GAPDH for TP53, SIRT1 and the phosphorylation levels of Histone 2A (pH2A.X), which is an established biomarker of DNA-damage^33^, were detected by Western blotting (fig. 3a) and full images of Western blots are shown in supplementary figure 3. All proteins were detected at the expected sizes of 110 kDa for SIRT1, 15 kDa for pH2A.X, 53 kDa for p53 and 38 kDa for GAPDH. While SIRT1 expression was not detectable in podocytes under control conditions, it significantly increased in podocytes incubated with healthy urine to 600% (p=0.01) and was even found to be further elevated when podocytes were incubated with AKI stage 2/3 urine to 1051% (p=0.01). In contrast, TP53 levels were found to be significantly downregulated by 85% when cells were incubated with healthy urine (p>0.05). When cells were incubated with the AKI stage 2/3 urine, TP53 expression was found to be significantly upregulated to 150% (p=0.05). Accordingly, the phosphorylation levels of H2A.X were found to be upregulated under both conditions, with healthy urine increasing the phosphorylation levels by 280% and the AKI 2/3 urine by 700%. Both changes were found to be significant (p<0.05). qRT-PCR analysis for TP53 and SIRT1 (fig.3b), revealed that mRNA expression for both is upregulated in podocytes treated with human urine. Healthy urine induced a 2.5-fold increase in TP53 mRNA expression and a 1.7-fold increase in TP52 expression, while AKI stage 2/3 urine induced an upregulation of TP53 mRNA by 3.5-fold and SIRT1 mRNA by 4.2-fold. Finally, we applied immunofluorescent-based detection of TP53 in cells exposed to AKI 2/3 urine (fig.3c), revealing an increase in positively stained cells.

**Figure 3:**
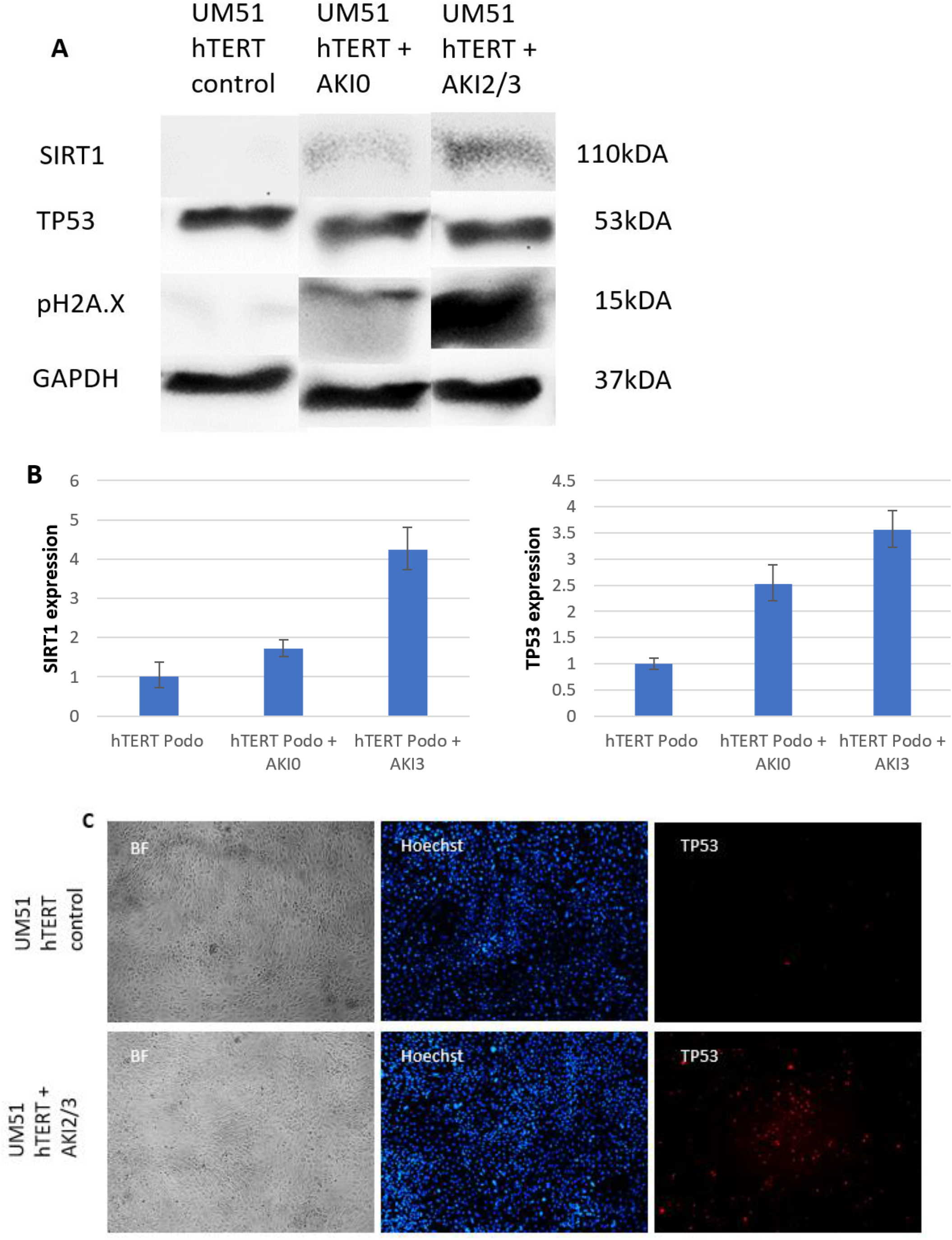
Upregulation of TP53 and SIRT1 induced by AKI stage 2/3 urine in podocytes. Podocytes were incubated with 72h post-surgery healthy and AKI stage 2/3 urine for 5 days. Relative protein expression normalized to GAPDH for SIRT1, TP53 and H2A.X phosphorylation was detected by Western blot (a). mRNA expression of *TP53* and *SIRT1* was determined by quantitative real time PCR (b). TP53 expression in Podocytes exposed to AKI 2/3 urine was detected by immunofluorescence-based staining (c).

**Figure 4:**
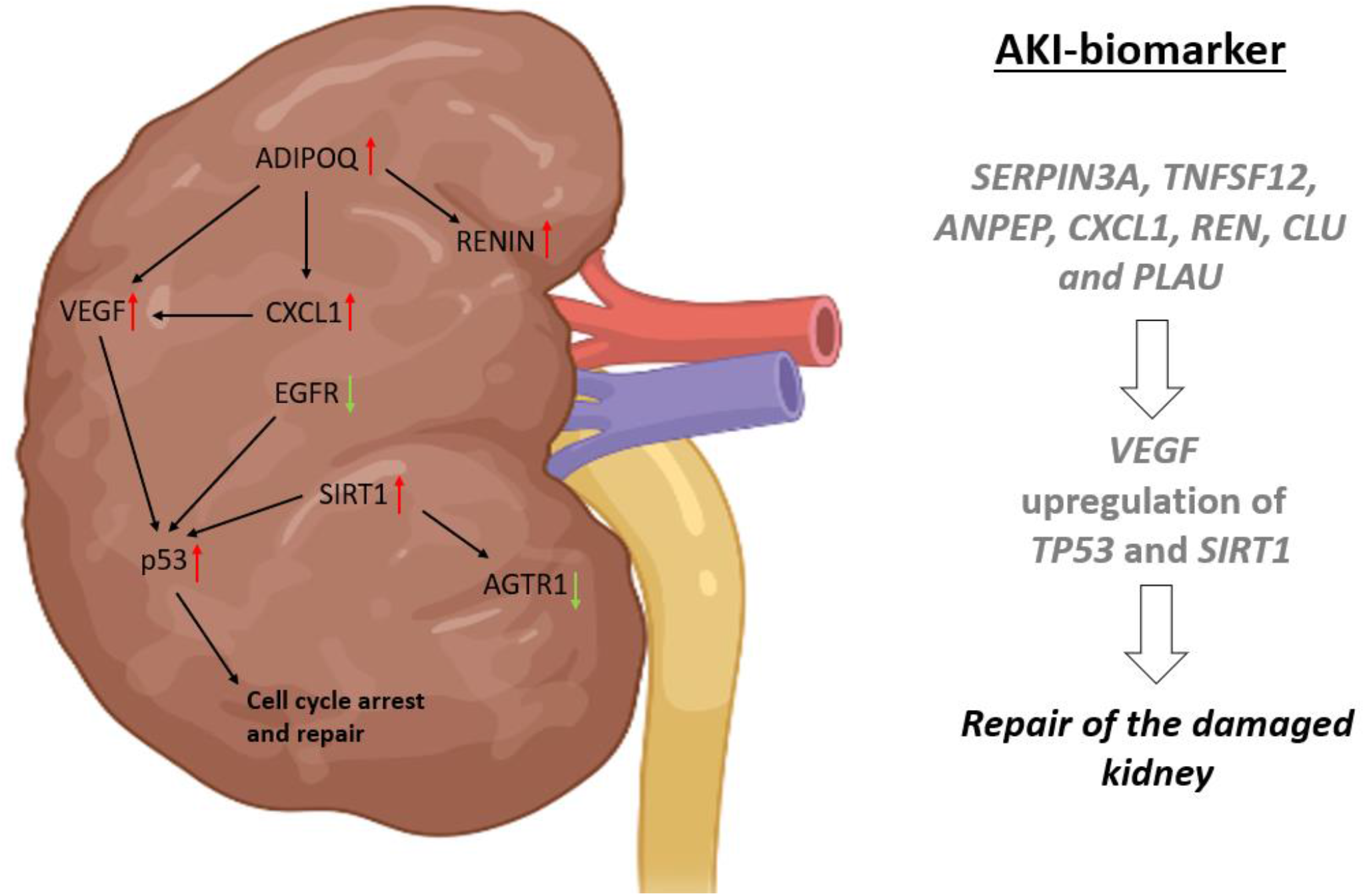
Proposed AKI signalling cascade. Urine-based biomarkers for the detection of AKI which are male-enriched (AHSG), female-enriched (CCN1, IL6, CCL2, THBS1, IL1RN) and common in both (VEGF, SERPIN3A, TNFSF12, ANPEP, CXCL1, REN, CLU and PLAU). Furthermore, mechanistically the cytokines ADIPOQ, EGF, EGFR, REN and VEGF present in the urine of AKI 2/3 patients trigger processes such as angiogenesis, needed to repair the damaged nephron and activation of TP53 and SIRT1 to maintain the balance between proliferation, angiogenesis, repair and cell cycle arrest. “Created with <u>BioRender.com.</u>“

## Discussion

In this study, by comparing urine-secreted cytokines of non-AKI and AKI 2/3-patients after cardiac surgery, we identified urine-based cytokine biomarkers of AKI. Since kidney disease can be difficult to identify in the clinic, because they do not cause any specific signs or symptoms^6^ and AKI can arise as part of other syndromes such as heart or liver failure and sepsis, variability between individuals and gender has to be accounted for. While male urine samples 24 h post-surgery revealed only seven cytokines as significantly altered, female urine samples contained a plethora of significantly altered cytokines. Common cytokines in both female and male urine samples provided 24h post-surgery were the upregulation of Adiponectin (ADIPOQ) and the downregulation of Epidermal growth factor (EGF) and Serpin Family A Member 3 (SERPIN3A). In contrast the analysis of urine from stage 2/3AKI patients 72 h post-surgery showed a dramatic increase in regulated cytokines. Here, male, and female samples showed an upregulation of Vascular Endothelial Growth Factor (VEGF), Alanyl Aminopeptidase (ANPEP), Chemokine (C-X-C Motif) Ligand 1 (CXCL1) and Renin (REN), while Clusterin (CLU), SERPIN3A, Plasminogen Activator Urokinase (PLAU) and Tumour necrosis factor ligand superfamily member 12 (TNFSF12) were found to be downregulated. Of note, the upregulation of ADIPOQ detected in24 post-surgery urine was reduced after 72 h. The same trend was observed for EGF in the female urine, while the downregulation was persistent in male AKI urine samples 72h post-surgery. Interestingly, a downregulation of secreted Interleukin 10 was only observed in male AKI urine samples 24 h and 72 h post-surgery. Based on these results we suggest, ADIPOQ, EGF and SERPIN3A as potential biomarkers, which might be able to detect AKI as early as 24 h post-surgery. For the later stages, common biomarkers for the detection of AKI in both male and female patients we propose, VEGF, SERPIN3A, TNFSF12, ANPEP, CXCL1, REN, CLU and PLAU. These markers in combination might present a robust strategy for identifying the development of AKI as early as 24 h or 72 h post-surgery. Many of these have already been described in the context of AKI^34–36^. The full list of significantly regulated cytokines is given in table 2.

**Table 2:** Summary of the identified biomarker for early diagnosis of AKI.

| Biomarker | Female 24 h post-surgery | Male 24 h post-surgery | Female 72 h post-surgery | Male 72 h post-surgery |
| --- | --- | --- | --- | --- |
| ADIPOQ | Up | Up |  |  |
| CCN1 | Up |  |  | Down |
| EGFR | Up |  |  | Up |
| IL6 | Up |  |  |  |
| CCL2 | Up | Down | Down | Up |
| THBS1 | Up |  |  |  |
| VEGF | Up |  | Up | Up |
| ANPEP | Down |  |  |  |
| B2M | Down |  | Down |  |
| CST3 | Down |  |  |  |
| DDP4 | Down |  |  |  |
| EGF | Down | Down | Down |  |
| IL1RN | Down |  | Up | Down |
| KLK3 | Down |  |  |  |
| SERPIN3A | Down | Down | Down | Down |
| TNFRSF1A | Down |  |  |  |
| TTF3 | Down |  |  |  |
| TNFSF12 | Down | Up | Down | Down |
| IL10 |  | Down |  | Down |
| TNFA |  | Down |  |  |
| ANPEP |  |  | Up | Up |
| CXCL1 |  |  | Up | Up |
| REN |  |  | Up | Up |
| CLU |  |  | Down | Down |
| MMP9 |  |  | Down |  |
| MME |  |  | Down |  |
| KLK3 |  |  | Down |  |
| AGER |  |  | Down |  |
| PLAU |  |  | Down | Down |
| AHSG |  |  |  | Up |
| ANXA5 |  |  |  | Down |
| IL6 |  |  |  | Down |
| SKP1 |  |  |  | Down |

At the molecular level, ADIPOQ might be of special interest since it has been described to directly regulate the expression three of the identified biomarkers which were upregulated 72 h post-surgery, namely CXCL1, VEGF and RENIN.

In cancer cells Adiponectin has been shown to induce CXCL1 secretion and thereby promoting tumour angiogenesis^27^. In addition CXCL1 has been reported to stimulate angiogenesis via the VEGF pathway^37,38^ and has even been suggested as a biomarker for patients suffering from colon cancer^39^. Furthermore, it has been shown that inhibition of the renin-angiotensin system results in significantly increased Adiponectin levels^40^. Therefore, we propose that the observed upregulation of secreted RENIN in AKI urine samples 72 h post-surgery might be causative for the downregulation of secreted ADIPOQ, while the early upregulation of secreted ADIPOQ results in the induction of the angiogenesis related genes CXCL1 and VEGF, thus reflecting the initiation of the repair processes in the damaged nephron. For instance VEGF has been shown to be an important mediator of the early and late phase of renal protective action after AKI in the context of stem cell treatment^35^.

EGF and EGFR both have been associated with AKI. Low levels of EGF have been reported to be predictive of AKI in newborns^41,42^. Of note, EGF has been reported to induce several matrix metalloproteinases^43^, and therefore the identified downregulation of EGF might be causative for the also observed downregulation of MME and MMP9 in female AKI urine 72 h post-surgery. EGFR has been reported to play an important role in renal recovery from AKI through PI3K-AKT-dependent YAP activation^44^. Indeed the EGFR promoter bears several p53-responsive elements^45^ and it has been reported that renal fibrosis is associated with EGFR and p53 activation^46^. Interestingly, the FDA approved test “NephroCheck®”, which is in clinical use and predicts the risk of developing moderate to severe AKI within 24h^47^, uses the following two biomarkers: tissue inhibitor of metalloproteinase 2 (TIMP-2) and insulin-like growth factor binding protein 7 (IGFBP7). Both are secreted by renal tubular cells in response to cellular stress and are involved in G1 phase cell cycle arrest^48^. This is achieved by the transcriptional activation of p27 via TIMP-2 and p53 and p21 via IGFBP7^49,50^. Further highlighting the involvement of TP53 activation in the clinical manifestation of AKI. Another study carried out by Montero et al., also documented that VEGF and p53 expression is significantly correlated and can be used as independent prognostic factors^28^. From our results, we conclude that secreted EGFR and VEGF are sufficient for upregulating p53 mRNA and protein levels in our immortalized podocyte cell line. Furthermore, another study carried out by Pierzchalski et al., showed that p53 is able to induce apoptosis via the activation of the Renin-Angiotensin System^29^. In two recent studies carried out in our laboratory, we demonstrated the devastating effect of the activated Renin-Angiotensin-System on podocyte morphology and functionality^51,52^. Therefore, we propose that the observed upregulated secretion of EGFR and VEGF mainly trigger angiogenesis and the activation of p53 and the Renin-Angiotensin system. This mode of action might be a plausible explanation for the progression from AKI to CKD. Considering the constant secretion of these factors, they should be investigated in urine samples from patients with AKI for more than three days after surgery and patients who progress to CKD. Interestingly, our recently published study investigating the secretome of urine from a CKD patient cohort from Ghana revealed the downregulation of secreted EGFR and VEGF ^53^. This makes it tempting to speculate that EGFR and VEGF might also be involved as initiators of AKI and downregulated in the progression to CKD. Especially VEGF in the context of CKD has been shown to have therapeutic potential by inducing renal recovery^54^. The exact mechanism underlying this protective effect of VEGF is unknown but kidney tissue after AKI is characterized by impaired endothelial proliferation and mesenchymal transition-both contributing to vascular refraction and subsequently to the progression to CKD^55^.

Finally, we found an upregulation of SIRT1 in podocytes incubated with the patient derived urine, which was found to be even more pronounced when AKI stage 2/3 urine was used. SIRT1 has been reported to be renal protective during AKI^23,56^. This protective effect might be caused by two distinct pathways which are regulated by SIRT1 activity. First, p53 is a direct substrate of the deacetylation activity of SIRT1 by regulating p53 activity, since p53 acetylation promotes the transactivation of numerous genes regulating cell cycle arrest, apoptosis and metabolic targets^57^. In this context it has been shown that SIRT1 overexpression promotes cellular survival in the presence of a cellular stressor^58,59^. The second pathway directly influenced by SIRT1 is the Renin-Angiotensin-Pathway^32^. In this context the overexpression of SIRT1 has been reported to attenuate Angiotensin II induced vascular remodelling and hypertension^60^, which might be caused by SIRT1-induced downregulation of the Angiotensin II type 1 receptor (AGTR1)^61^. SIRT1 has been reported to increase lifespan in lower organisms, such as *Caenorhabditis elegans*^*62*^ and *Drosophila melanogaster*^*63*^, and recently our own group reported the age-associated decline in SIRT1 mRNA and protein in Urine derived Renal Progenitor cells^64^. Since AKI has been discussed as a condition of renal senescence^65^, the expression level of SIRT1 seems to be of major importance for the prevention of AKI.

In summary, we propose putative urine-based biomarkers (24 and 72 h) for the detection of AKI which are male-enriched (AHSG), female-enriched (CCN1, IL6, CCL2, THBS1, IL1RN) and common in both (VEGF, SERPIN3A, TNFSF12, ANPEP, CXCL1, REN, CLU and PLAU). Furthermore, mechanistically the cytokines ADIPOQ, EGF, EGFR, REN and VEGF present in the urine of AKI 2/3 patients trigger processes such as angiogenesis, needed to repair the damaged nephron and activation of p53 and SIRT1 to maintain the balance between proliferation, angiogenesis, repair and cell cycle arrest. On this basis, we propose a specific signalling cascade induced by damage to the kidney resulting in the establishment and propagation of AKI (fig.4). Furthermore, the Renin-Angiotensin pathway might have major implications.

## Material and Methods

### Study design

After approval of the local Ethics committee of Heinrich-Heine University Duesseldorf, Germany (local trial number 5803, clinicaltrials.gov: NCT03089242), adult patients scheduled for cardiac surgery were enrolled after they had provided written informed consent to participate in the study.

Urinary samples of 6 patients with moderate to severe AKI (AKI stage 2 or 3) and of 6 matched controls (heart surgery, no AKI post-operatively) were centrifuged (3500 rpm, 5 minutes) and stored until further analysis.

### Secretome analyses

Urine samples were analyzed using the Proteome Profiler Human Kidney Biomarker Array Kit (#ARY019) distributed by Research and Diagnostic Systems, Inc. (Minneapolis, Minnesota, United States) as described by the manufacturer and in our previous publication^53^. In brief, membranes were blocked with the provided blocking buffer and each urine sample was incubated with the 15μL of reconstituted Detection Antibody Cocktail for 1h at Room temperature. After removing the blocking buffer, the urine samples were incubated on the membranes at 4 °C overnight on a rocking platform. After three consecutive washing steps each for 10min at room temperature on a rocking platform, the membranes were incubated with the diluted Streptavidin-HRP antibody mix. After three consecutive washing steps each for 10min at room temperature on a rocking platform, fluorescence signals were visualized with enhanced luminescence (WesternBright Quantum, Advansta, San Jose, United States).

Obtained images were analyzed by using the Image J software^66^ with the Microarray Profile plugin by Bob Dougherty and Wayne Rasband (https://www.optinav.info/MicroArray_Profile.html). The integrated density generated by the Microarray profile plugin function Measure RT was used for follow-up processing which was performed in the R/Bioconductor environment^67^. Arrays were normalized employing the Robust Spline Normalization from the Bioconductor lumi package^68^. A threshold for background intensities was defined at 5% of the range between maximum and minimum intensity and a detection-p-value was calculated according to the method described in Graffmann et al.^69^.

### Cell culture conditions

The cell line UM51-hTERT was derived and cultured as described in^5126^. For the experiments, non-AKI and AKI 2/3 patient urine samples 72h post-surgery were pooled and mixed with the culture medium to a final concentration of 10%. Cells were incubated with this medium for five days.

### Relative Quantification of podocyte-associated gene expression by real-time PCR

Real-time PCR of podocyte-associated gene expression was performed as follows:

Real time measurements were carried out on the Step One Plus Real-Time PCR Systems using MicroAmp Fast optical 384 Well Reaction Plate and Power Sybr Green PCR Master Mix (Applied Biosystems, Foster City, United States). The amplification conditions were denaturation at 95 °C for 13 min. followed by 37 cycles of 95 °C for 50 s, 60 °C for 45 s and 72 °C for 30 s. Primer sequences are listed in supplementary table 3.

### Immunofluorescence-based detection of protein expression

Cells were fixed with 4% paraformaldehyde (PFA) (Polysciences, Warrington, United States). Unspecific binding sites were blocked by incubation with blocking buffer, containing 10% normal goat or donkey serum, 1% BSA, 0.5% Triton, and 0.05% Tween, for 2 h at room temperature. Incubation of the primary antibody was performed at 4 °C overnight in staining buffer (blocking buffer diluted 1:1 with PBS). After at least 16 h of incubation, the cells were washed three times with PBS/0.05% Tween and incubated with a 1:500 dilution of secondary antibodies. After three additional washing steps with PBS/0.05% Tween the cell and nuclei were stained with Hoechst 1:5000 (Thermo Fisher Scientific, Waltham, United States). Images were captured using a fluorescence microscope (LSM700; Zeiss, Oberkochen, Germany) with Zenblue software (Zeiss). Individual channel images were processed and merged with Fiji. Detailed Information of the used antibodies are given in supplementary table 4.

### Western Blotting

Protein isolation was performed by lysis of the cells in RIPA buffer (Sigma-Aldrich, St. Louis, United States), supplemented with complete protease and phosphatase inhibitors (Roche, Basel, Switzerland). Proteins were separated on a 7.5% Bis-Tris gel and blotted onto a 0.45 μm nitrocellulose membrane (GE Healthcare Life Sciences, Chalfont St. Giles, United Kingdom). After blocking the membranes with 5% skimmed milk in Tris-buffered Saline Tween (TBS-T), they were incubated overnight with the respective primary antibodies (supplementary table 3). Membranes were washed 3x for 10 min with TBS-T. Secondary antibody incubation was performed for 1h at RT and membranes were subsequently washed 3x for 10 min with TBS-T. Amersham ECL Prime Western Blotting Detection Reagent was used for the chemiluminescent detection (GE Healthcare Life Sciences) and captured with the imaging device Fusion FX.

## Notes

### Competing Interest Statement

The authors have declared no competing interest.

