## supplementary files for "Cytokines in the Urine of AKI patients regulate TP53 and SIRT1 and can be used as biomarkers for the early detection of AKI"

Supplemental Material Table of Contents

Table 3: PCR Primers

Table 4: antibodies

Supplementary figure 1: Cytokine Array pictures 24 post-surgery.

Supplementary figure 2: Cytokine Array pictures 72 post-surgery.

Supplementary figure 3: Quantification of the western blots and full-sized blot images.

Table 1: Primer

| Primer name | Sequence | Annealing temperature (°C) | Product length (bp) |
|---|---|---|---|
| RPL0 s | 5‘-TCGACAATGGCAGCATCTAC-3‘ | 60 | 195 |
| RPL0 as | 5‘-ATCCGTCTCCACAGACAAGG-3‘ | | |
| SIRT1 (Exon 1-2) s | 5‘-AGGGCGAGGAGGAGGAAGAG-3‘ | 60 | 122 |
| SIRT1 (Exon 1-2) as | 5‘-GGCTCTATCCTCCTCATCACTTTC-3‘ | | |
| TP53 s | 5‘-CAGGGCAGCTACGGTTTCC-3‘ | 60 | 157 |
| TP53 as | 5‘-CAGTTGGCAAAACATCTTGTTGAG-3‘ | | |

Table 2: antibodies

| Antigen | Company | Dilution (IF/WB) |
|---|---|---|
| Rabbit GAPDH | Cell Signaling #2118 | n.a./1:1000 |
| Rabbit pH2A.X (Ser139) | Cell Signaling #9718S | 1:200/1:1000 |
| Mouse TP53 | Merck Millipore #OP43 | n.a./1:1000 |
| Mouse SIRT1 | Abcam #ab110304 | 1:200/1:1000 |
| Anti-mouse Alexa Fluor 488 | Invitrogen #A11070 | 1:500 |
| Anti-mouse HRP-labeled | Thermo Fisher Scientific #NA931 | 1:4000 |
| Anti-rabbit HRP-labeled | Cell Signaling #7074S | 1:1000 |

AKI stage 2/3

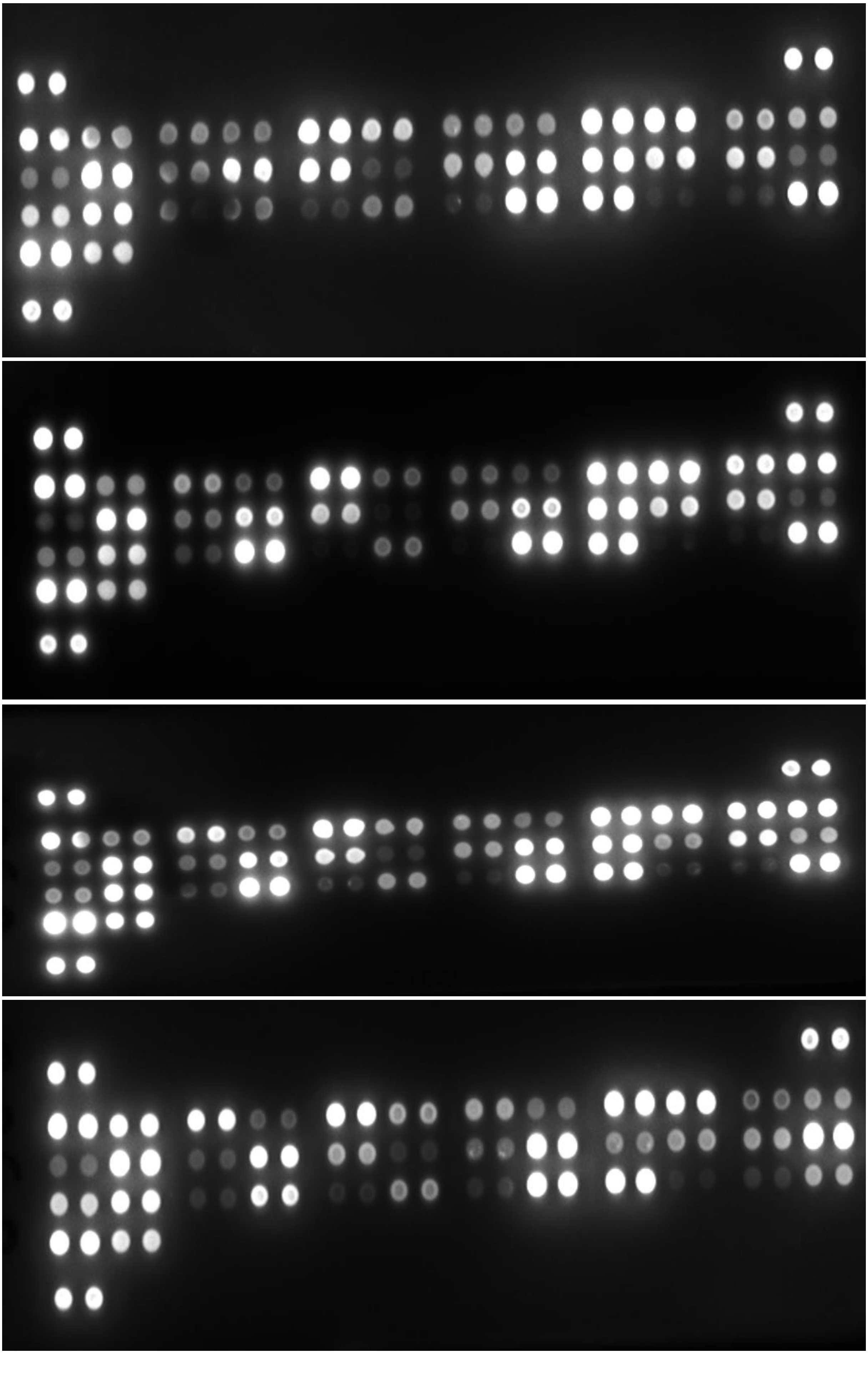

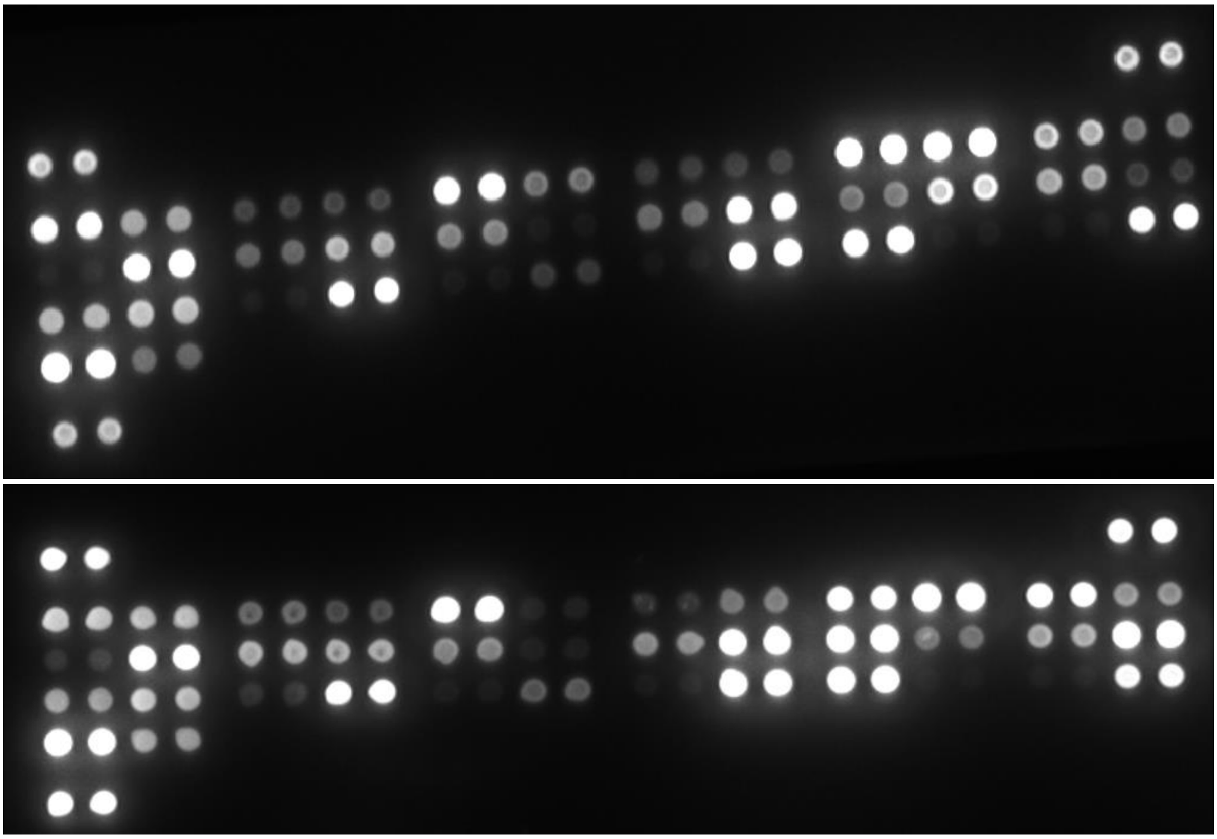

healthy

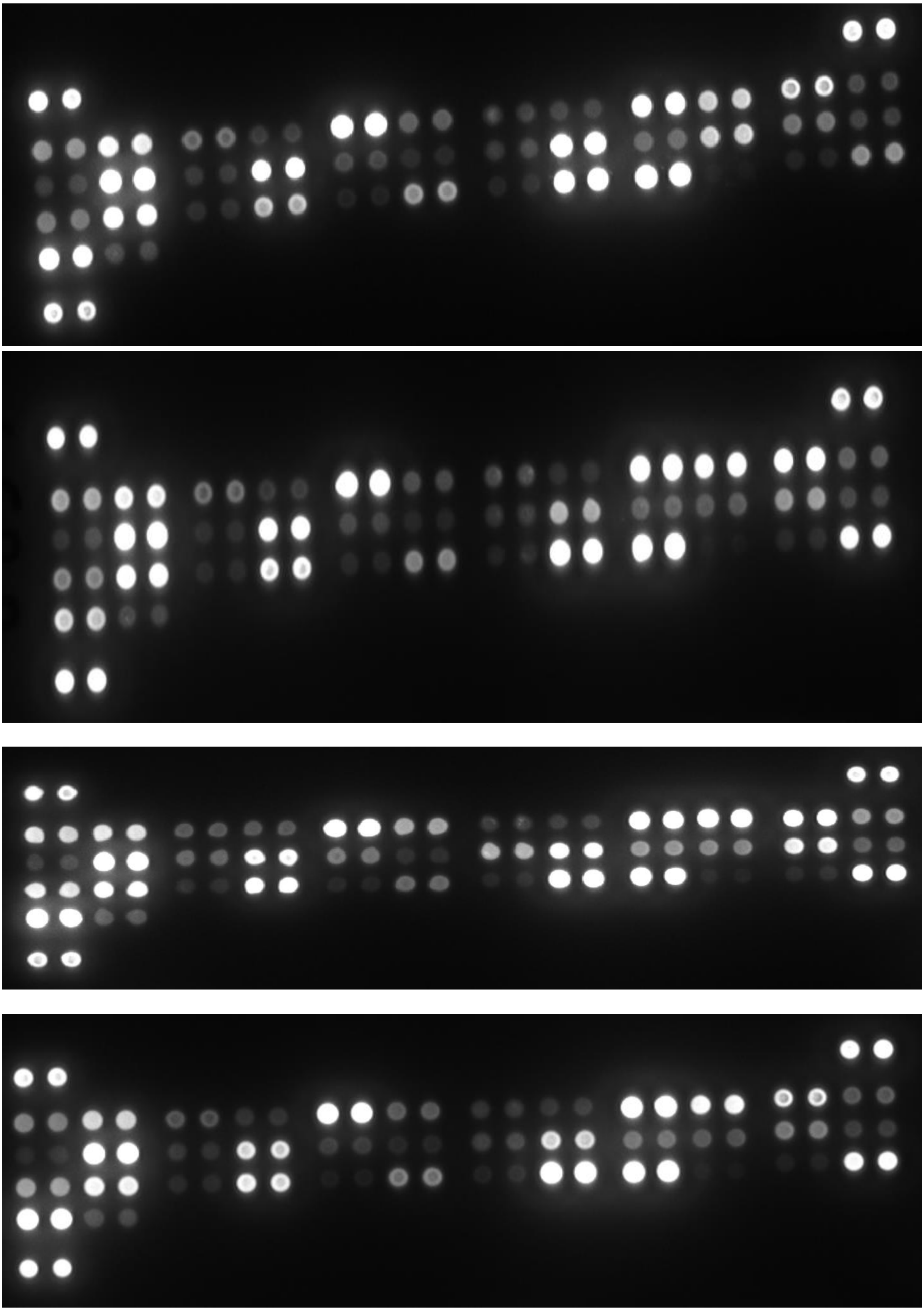

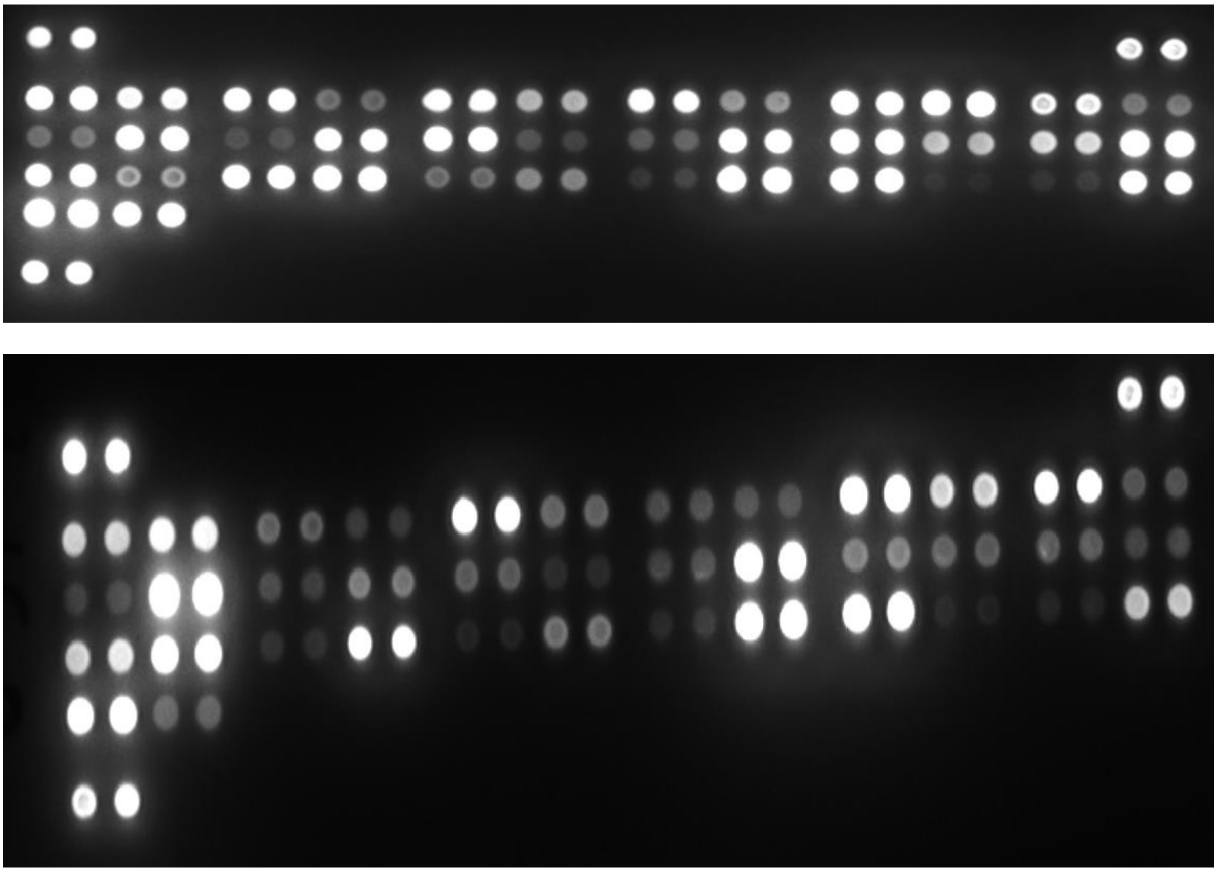

**Supplementary figure 1: Cytokine Array pictures 24 post-surgery.**

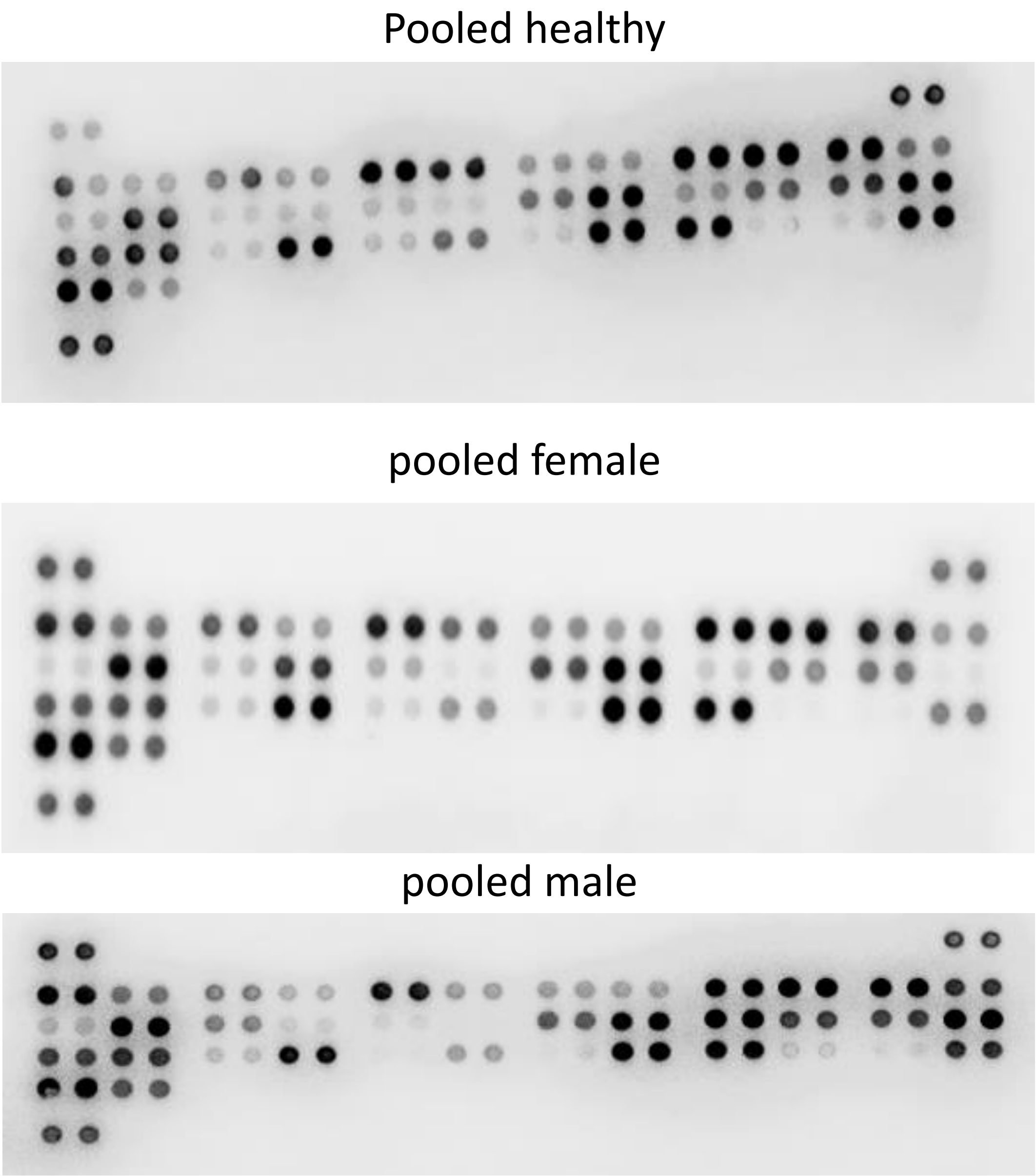


**Supplementary figure 2: Cytokine Array pictures 72 post-surgery.**

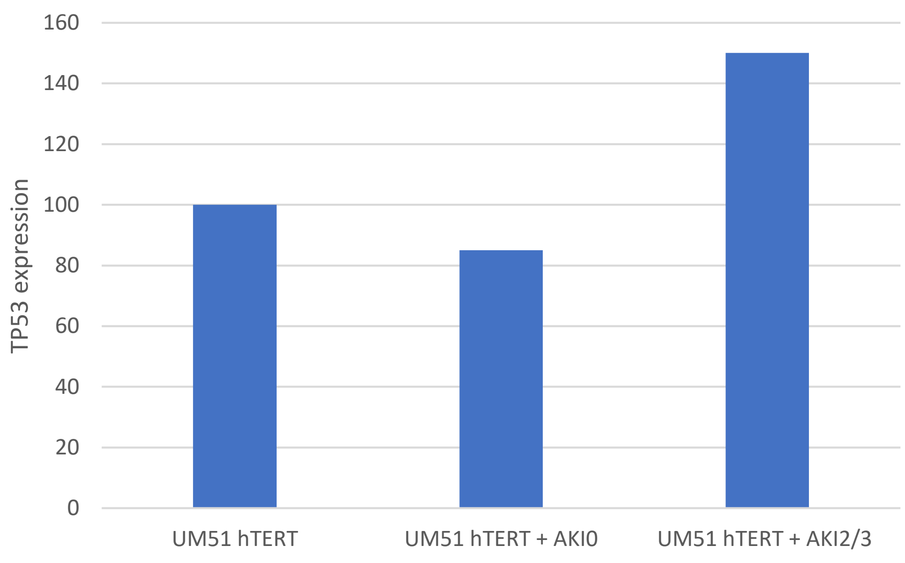
160
140
120
100
80
60
40
20
0
TP53 expression
UM51 hTERT
UM51 hTERT + AKI0
UM51 hTERT + AKI2/3

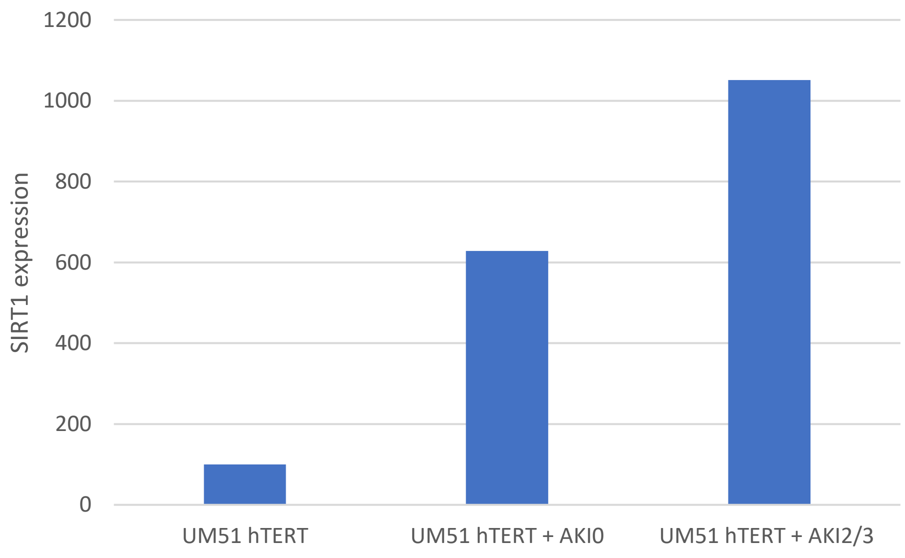
1200
1000
800
600
400
200
0
SIRT1 expression
UM51 hTERT
UM51 hTERT + AKI0
UM51 hTERT + AKI2/3

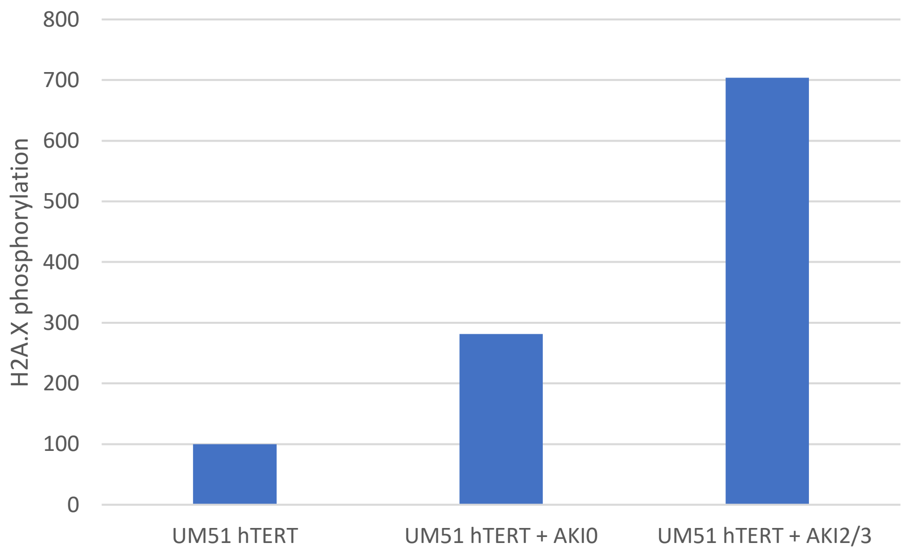
800
700
600
500
400
300
200
100
0
H2A.X phosphorylation
UM51 hTERT
UM51 hTERT + AKI0
UM51 hTERT + AKI2/3

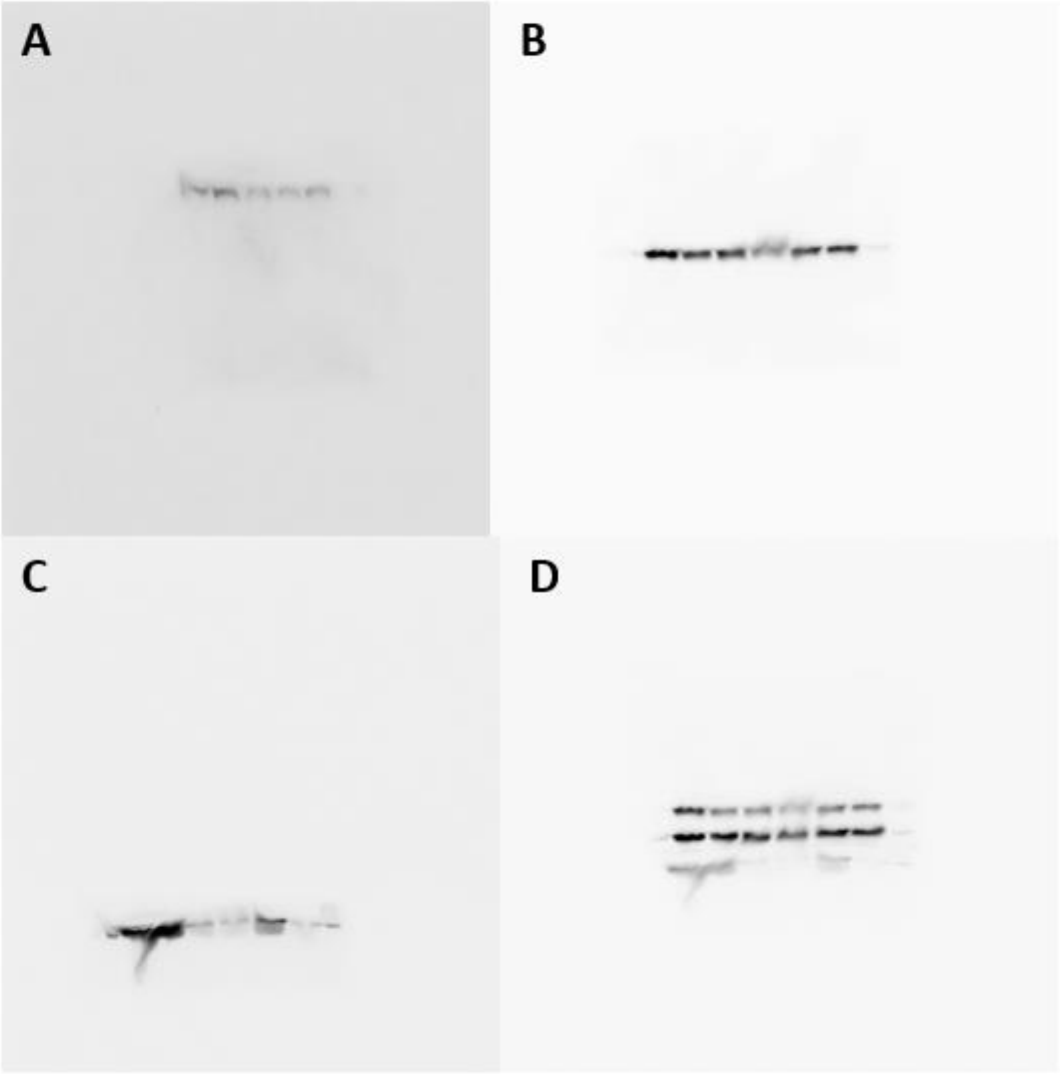


**Supplementary figure 3: Quantification of the western blots and full-sized blot images.**
Western blot full-sized images are given for SIRT1 (A), TP53 (B), H2A.X (C) and GAPDH (D). UM51-hTERT control podocytes, cultured with urine from AKI0 patients and with urine from AKI 2/3 patients are represented in lane 1, 2 and 5, respectively.
